# Offline Reconstruction of Diffusion MRI Acquisitions for Comparison Between Complex PCA-based and AI-based Denoising

**DOI:** 10.1101/2025.11.07.682283

**Authors:** Francesco D’Antonio, Shaun Warrington, Jose-Pedro Manzano-Patron, Paul S. Morgan, Stamatios N. Sotiropoulos

## Abstract

**Purpose:** Optimal diffusion MRI (dMRI) data for image denoising is often unavailable from scanner reconstruction. In this work, we make available an offline reconstruction pipeline for GE dMRI acquisitions, giving access to complex dMRI data. Furthermore, we compare the efficacy of GE HealthCare’s AIR-Recon DL™ (ARDL), a proprietary convolutional neural network-based reconstruction and denoising approach, to open-source PCA-based MPPCA_SVS_ and NORDIC denoising methods on high-resolution dMRI data.

**Methods:** We developed an end-to-end offline dMRI reconstruction pipeline for GE HealthCare acquisitions, augmenting the Orchestra software development kit, and validated its output against scanner reconstruction. We used it to compare MPPCA_SVS_, NORDIC and ARDL denoising approaches, considering underlying metrics reflecting noise variance and bias, such as the ADC profiles in highly anisotropic areas, and downstream measurements, such as fiber orientation estimation and white matter tractography.

**Results:** Our validated offline reconstruction supports various in-plane/out-of-plane accelerations and partial Fourier reconstruction methods. Unlike scanner reconstruction, it provides access to complex dMRI data, enabling denoising in the complex domain, which demonstrated superior noise floor suppression compared to magnitude-constrained denoising. PCA-based denoising methods had improved spatial resolution, contrast-to-noise and more robust fiber orientation estimation compared to ARDL.

**Conclusion:** We found significant gains in dMRI data quality when using the proposed offline reconstruction pipeline, allowing complex-domain denoising to obtain high-quality data at high spatial resolution and b-value, using a wide-bore scanner and a standard PGSE EPI sequence. MPPCA_SVS_ and NORDIC (4D PCA-based) outperformed ARDL (2D) in terms of spatial resolution, reduction of noise-floor bias and variance.

## 1. Introduction

The value of denoising diffusion MRI (dMRI) data has been demonstrated in various scenarios, enhancing fidelity of dMRI-derived microstructural estimates [1–3], increasing spatial resolution [2, 4], and improving consistency between scanners [5]. DMRI contrast is derived from signal attenuation, leading to low SNR, especially when increasing resolution or diffusion weighting. This results in low signals being rectified to the noise floor [6], leading to biases in parametric maps and downstream fiber modelling [3, 7–9]. To overcome these challenges, several denoising approaches have been developed using PCA-based, low-rank reduction of signals [5, 10–17]. These methods exploit the spatial, angular and diffusion-weighted redundancy in dMRI signals to estimate and remove noise-dominant principal components from local patches. A breakthrough in the field occurred when Random Matrix Theory was used to derive data-driven estimates for the threshold between noise and signal-dominant principal components [12, 13]. This is achieved by fitting the Marchenko-Pastur (MP) distribution [18] to the distribution of principal components, eliminating the need for a separate estimate of the noise variance [14, 19]. Refinements for threshold estimation, which account for signal-dominant components that alter the theoretical MP distribution, have been implemented to better estimate the signal-noise boundary [10, 16, 20]. Additionally, to estimate noise-free signal more accurately, optimal shrinkage of principal components is used to mitigate eigenvalue inflation due to noise [21] using nuclear [1] or Frobenius norms [5, 16]. Regardless of the exact implementation, it has been demonstrated that the domain (magnitude/complex) into which dMRI denoising is applied can have significant implications for subsequent analysis, affecting noise-induced variance, bias, and true resolution in different ways [1, 2, 5, 22]. A challenge, however, is that across scanner vendors it is not straightforward (or not feasible at all in some cases) to control reconstruction parameters, such as apodization filters, or even access complex data, which would improve denoising outcomes.

In addition to these PCA-based approaches, scanner manufacturers have recently introduced AI-driven reconstruction and denoising solutions to enhance resolution and SNR while maintaining or reducing scan time [23–27]. These methods are integrated into the scanner reconstruction process, making them easily implementable in clinical protocols, and benefit from having access to unfiltered, complex domain data for optimal noise reduction. One such method is AIR™ Recon DL (ARDL) from GE HealthCare (Milwaukee, USA), a reconstruction and denoising convolutional neural network trained on millions of MRI images from various scan locations and acquisition methods, which is used to and remove thermal noise and Gibbs ringing artefacts [23]. ARDL operates on 2-dimensional image slices in the complex domain. The trade-off between resolution, noise, and scan time typically governs acquisition choices for MRI, particularly for noise-prone modalities, such as dMRI, making it an ideal modality for assessing the performance of this denoising method. However, it is unclear how DL-based approaches, such as ARDL, perform when compared against PCA-based approaches.

In this paper, we address the above questions. Firstly, we develop an end-to-end offline reconstruction pipeline compatible with GE dMRI acquisitions. It takes raw, single-channel, complex k-space data as input and provides access to complex dMRI data. The pipeline supports retrospective switching of filters and partial Fourier reconstruction methods and is compatible with in-plane and out-of-plane accelerated acquisitions. This is necessary as scanner reconstruction on GE scanners does not provide access to complex dMRI data. Secondly, we demonstrate the application of this offline reconstruction pipeline by comparing PCA-based denoising in the complex domain with GE’s proprietary ARDL on standard PGSE acquisition from a wide-bore scanner.

Specifically, we begin by validating the pipeline by comparing the scanner’s and the pipeline’s magnitude reconstruction across different acceleration and reconstruction parameters. We then compare ARDL with two commonly used denoising methods, NORDIC [11] and MPPCA_SVS_ [5, 12, 13] (MPPCA with singular value shrinkage), as they can be easily implemented in the complex domain. We also include MPPCA_SVS_ denoising in the magnitude domain to highlight improvements in denoising outcomes when using the offline reconstruction pipeline’s complex-domain data, over magnitude-constrained denoising available from native scanner reconstruction. To evaluate denoising outcomes, we consider quantitative metrics of noise variance, noise floor bias and proxies of noise distribution within the brain. We also evaluate the downstream effects of denoising through fiber orientation estimation and white matter bundle segmentation with probabilistic tractography.

## 2. Methods

### 2.1 Image Reconstruction Pipeline

We developed a comprehensive and scalable offline reconstruction pipeline for dMRI acquisitions on GE HealthCare scanners, starting from single-channel k-space data from individual coil elements and producing coil-combined images in both complex and magnitude domains. The pipeline can modify several reconstruction steps and handle multiple accelerated acquisition options.

We used Python (version 3.12.6) and built upon the C++ Orchestra software development kit (SDK), which provides simple example reconstruction scripts and is available through GE HealthCare’s WeConnect website (https://weconnect.gehealthcare.com/) with the appropriate research license. The pipeline takes as input ScanArchive.h5 files (k-space, individual coil data) and outputs DICOMs and NIfTI images (either magnitude or complex) as shown in Figure 1. Complementing further the capabilities of the default Orchestra SDK scripts, our workflow allows:

**Fig. 1.**
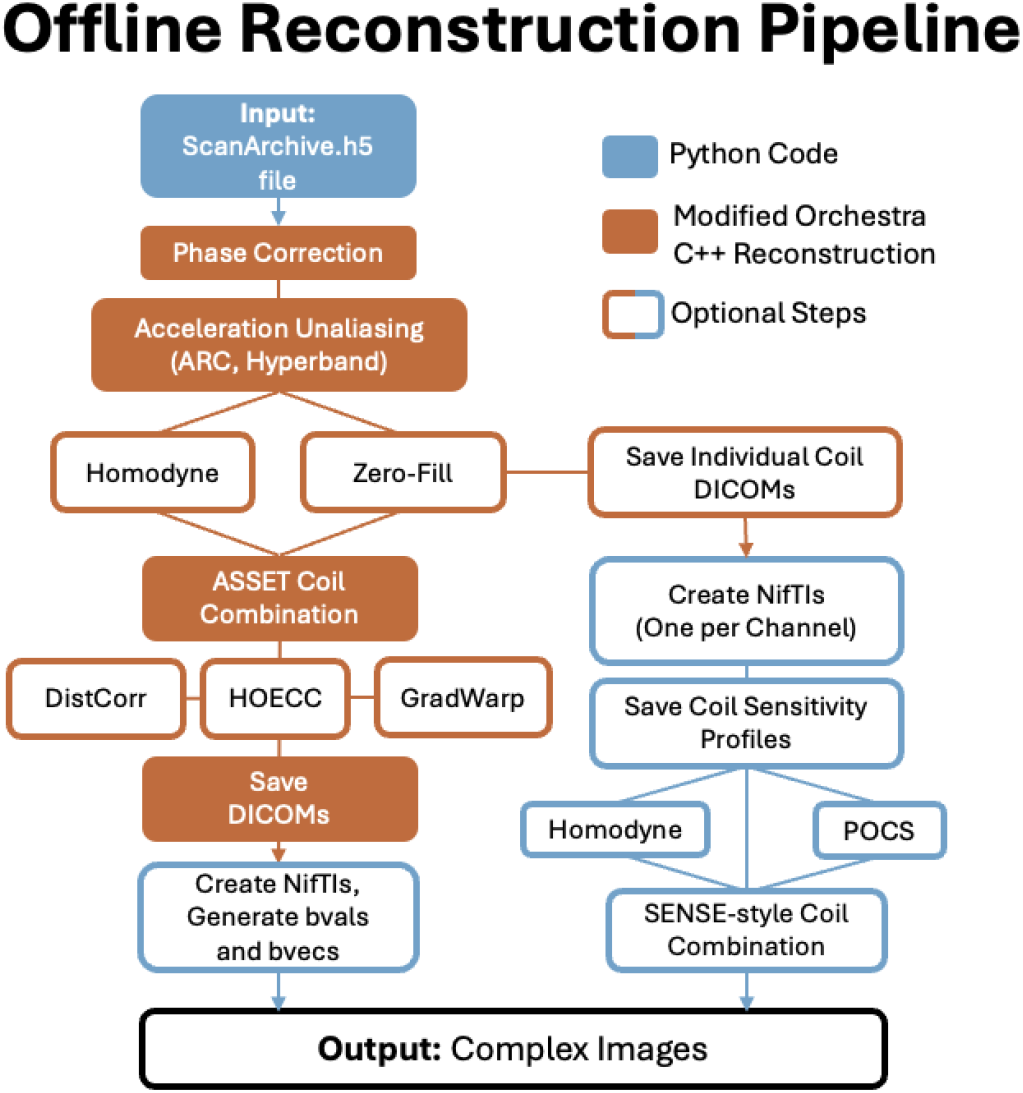
Outline of the offline reconstruction pipeline. Orange steps are modified C++ Orchestra SDK code; in blue are additional steps using custom Python code. Boxes with only colored borders (filled in white) are optional steps. The pipeline is implemented within a container for reproducibility and ease of use across operating systems.

i. Reconstruct channel-combined complex (magnitude/phase or real-rotated) data.
ii. Support of in-plane (ASSET-SENSE and ARC-GRAPPA) and out-of-plane accelerations, with proper reconstruction of HyperBand acquisitions for simultaneous multi-slice (SMS) accelerations.
iii. Retrospective reconstruction of partial Fourier acquisitions, using zero-filling, POCS [28] and Homodyne [29] methods.
iv. Access to single-channel complex images and extraction of coil sensitivity profiles to be used for sensitivity-weighted channel combination.
v. Proper indexing of dMRI volumes and slices for all combinations of in-plane (SENSE-like, GRAPPA-like) and out-of-plane (SMS) accelerations.
vi. Optional flags to modify several reconstruction steps retrospectively, including switching on/off gradient non-linearity correction, on/off apodization filters, on/off high-order eddy current correction and switching between complex (zero-filling) and homodyne reconstruction [29] for partial Fourier acquisitions.
vii. Accounting for and preserving phase encoding direction and correct creation of diffusion b values (bval) and diffusion-weighted direction vector (bvec) files for subsequent spin-echo fieldmap-based corrections of susceptibility-induced distortions.

To enable the reproducible and straightforward implementation of the pipeline across various operating systems and high-performance clusters, the offline reconstruction is embedded into Docker (https://docs.docker.com/) or Singularity/Apptainer (https://apptainer.org/) containers. Additionally, the C++ source code for the pipeline is available for customisation. The pipeline and source code is available through GE HealthCare’s WeConnect website with the appropriate research license.

To evaluate the capabilities of the pipeline, data using SENSE-like acceleration (ASSET factors 1 and 2), GRAPPA-like acceleration [30] (ARC factors 1 and 2), simulataneous multi-slice [31–33] (HyperBand factors 1 and 4), and partial Fourier reconstruction methods (homodyne or zero-filled), were acquired and reconstructed offline, using the developed reconstruction pipeline, as well as on the scanner. The magnitude reconstruction from both approaches was compared to validate the offline pipeline’s output.

### 2.2 Denoising Comparisons

The offline reconstruction was developed with denoising in mind and hence offers the ability to perform denoising either in magnitude or complex domain at various stages of the reconstruction, for instance, prior to or after channel combination. In this work, we considered two PCA-based denoising methods applied after channel combination:

- **MPPCA**_**SVS**_: using 7×7×7 patches (using the smallest odd N such that NxNxN > number of volumes acquired, using the heuristic in MRTrix [34]), using eigenvalue threshold estimation from Cordero-Grande et al. [10], and singular value shrinkage using Frobenius norm [21], as implemented in the DESIGNER pipeline (v2.0.11) [35, 36]. This was applied to both complex (real rotated homodyne [29]) data (MPPCA*_SVS_) and magnitude (|MPPCA|_SVS_) data.
- **NORDIC:** using 13×13×13 patches (so that the number of voxels in a patch and the number of diffusion volumes are approximately in a ratio of 11:1, as suggested in the NORDIC code) (https://github.com/SteenMoeller/NORDIC_Raw) [11] applied to complex (real rotated homodyne) data (NORDIC*). We did not provide noise only volumes, therefore the NORDIC method normalizes the data using an estimated g-factor map using 2D MPPCA [12, 13] noise residuals, then calculates low rank thresholds using generated patch-seized noise matrices.

We subsequently used the reconstruction pipeline to compare these PCA-based denoising approaches with GE’s AI-based ARDL. Starting from raw k-space data, we retained the real-channel rotated data from homodyne reconstruction [29] (referred to as complex data in the rest of the paper) and used this for denoising in the complex domain. ARDL, which also operates in the complex domain, was applied with the “high” denoising weighting hyperparameter (from low, medium, and high options from the scanner console interface) on the same data, using retrospective reconstruction on the scanner. As ARDL denoising requires a fixed-size 256×256 input matrix by design, k-space was manipulated by placing the acquired k-space in a zero-filled 256×256 matrix during reconstruction. After ARDL was applied, data were downsampled to the acquisition matrix size using spline interpolation [37, 38]. An overview of the reconstruction and processing pipeline is shown in Figure 2.

**Fig. 2.**
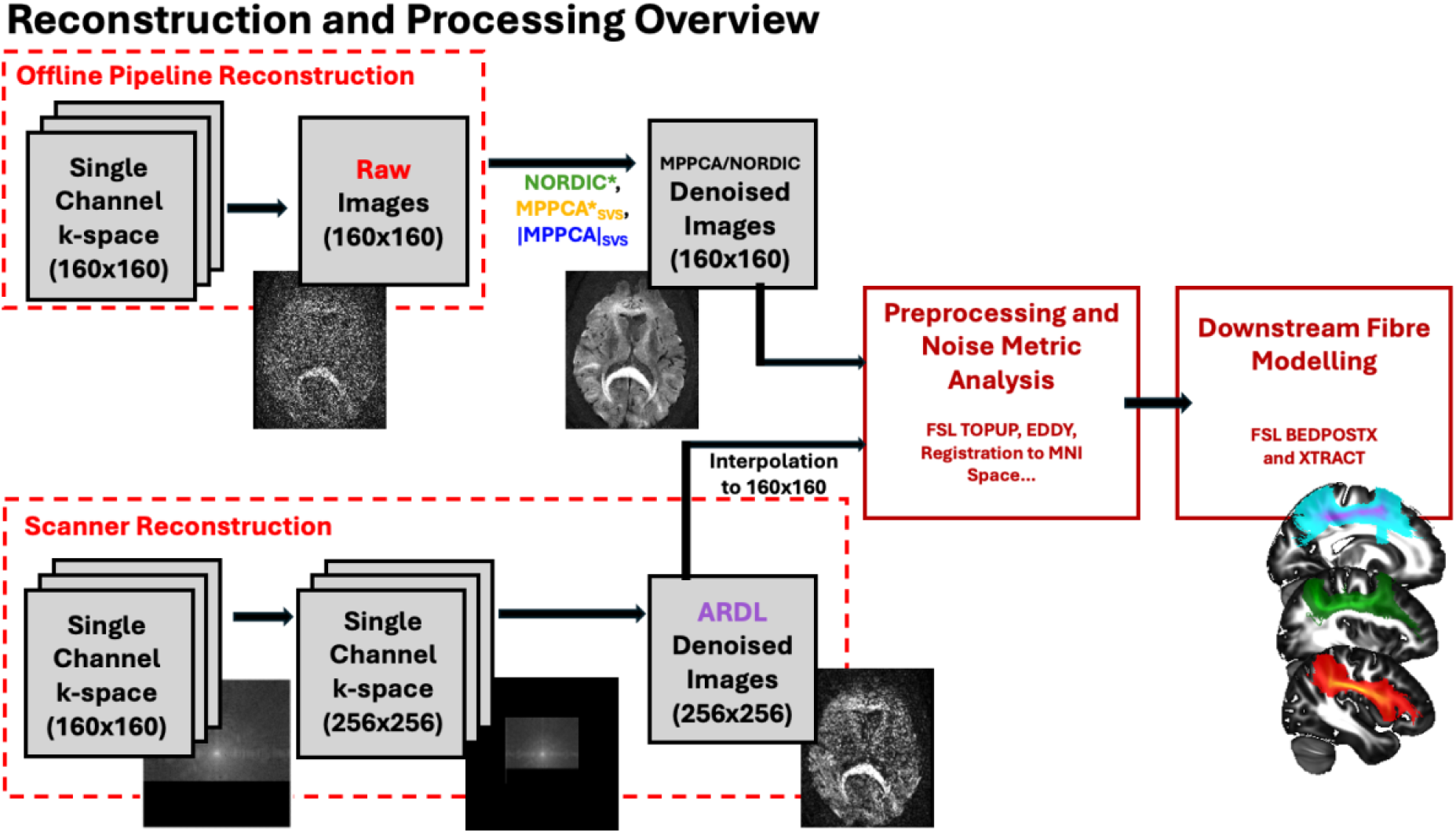
Overview of processing steps for comparison across denoising methods. For ARDL reconstruction and denoising, the k-space is resampled to a 256×256 matrix and later (post-ARDL) downsampled to the native matrix size.

### 2.2 Data Acquisition

Data for offline reconstruction pipeline validation were acquired on a GE HealthCare 3T Premier wide-bore scanner (Software Version MR29.2; GE HealthCare, Milwaukee, USA) at the University of Nottingham. Data from a single participant was acquired using 2 mm isotropic, 6 directions, b=1000 s/mm^2^ PGSE-EPI at various acceleration parameters: no acceleration (TR/TE = 8.0s/57.2ms), ASSET=2 (TR/TE = 8.0s/57.3ms), ARC=2 HyperBand=4 (TR/TE = 8.0s/59.7ms) as well as a high-resolution 1.3 mm isotropic HCP-style data [39, 40], (98 volumes/shell, two shells: b=1500, 3000 s/mm^2^, HyperBand=4, TR/TE=4.8s/88.3ms).

Data for denoising comparisons were acquired at the Cambridge University Hospitals NHS Foundation Trust (Cambridge, UK) for four healthy participants (two female, age range 27 to 54) on a GE HealthCare 3T Premier wide-bore scanner with a 48-channel head coil (Software Version MR30.1; GE HealthCare, Milwaukee, USA). Data were acquired using a PGSE-EPI sequence following the acquisition protocols used in the Human Connectome Project (HCP) [39, 40], with 1.25 mm isotropic voxel size, 98 volumes/shell, two shells: b=1500, 3000 s/mm^2^, and 14 b=0 s/mm^2^ volumes throughout the acquisition with one reverse gradient (AP/PA) b=0 s/mm^2^ acquired prior to the main data, HyperBand=4, TR/TE=4.8s/88.3ms, 60% partial Fourier with Homodyne reconstruction. A 1 mm isotropic T1-weighted MPRAGE with higher order shimming was acquired to create white matter masks and improve registration to MNI templates.

All data were acquired after informed consent was obtained from all participants. Ethical approval was respectively obtained from the University of Nottingham Faculty of Medicine and Health Sciences Research Ethics Committee (B12012012a_073) and from the East of England regional review board, Cambridgeshire and Hertfordshire Research Ethics Committee (08-H0311-117).

### 2.3 Pre-Processing and Analysis

Post-denoising, identical distortion correction across denoising methods was carried out using a standard FSL-eddy pipeline (FSL v6.0.6.4) [41, 42] on the images, including susceptibility-induced, eddy-current distortion and motion correction. We evaluated several aspects of denoising impact, from raw data to downstream analysis, as recommended in Manzano-Patron et al. [2]. To assess denoising efficacy with respect to variance suppression, SNR and angular CNR were obtained using FSL-eddyQC [43]. To assess efficacy with respect to bias and noise floor suppression, we examined the distribution, dynamic range and ADC profile of the signal, before and after denoising. To account for noise spatial non-stationarities, noise distributions were estimated from inside the brain (rather than background) using maximally attenuated (i.e. high b-value) CSF signal from the ventricles [44]. Noise-floor-induced signal rectification was assessed by examining the signal attenuation and ADC measurements of parallel vs perpendicular (i.e. maximally vs minimally attenuated) to principal fiber directions (DTI V_1_) in highly anisotropic voxels [44], in the corpus callosum, using single-subject warped masks. Downstream effects of denoising methods were also assessed through fiber orientation estimation and probabilistic tractography, carried out using FSL’s bedpostX [45, 46] and XTRACT [47] for each denoising mode and we compared session-wise tract density maps with average probabilistic tract density maps from a 50-subject HCP average [47].

## 3. Results

### 3.1 Offline Reconstruction Features and Validation

We validated the offline reconstruction by comparing magnitude images between the scanner and the offline reconstruction pipeline, as complex dMRI data are not available from the scanner reconstruction. We performed comparisons across resolutions and b-values, as shown in Figure 3, for various in-plane (no acceleration, ASSET=2, and ARC=2) and out-of-plane (no HyperBand and HyperBand=4) acceleration. In all cases, the offline magnitude reconstruction matched the scanner reconstruction almost perfectly when no SMS acceleration was used. When SMS acceleration was used minor signal differences were seen within the brain, but these differences were below 1% in absolute value.

**Fig. 3.**
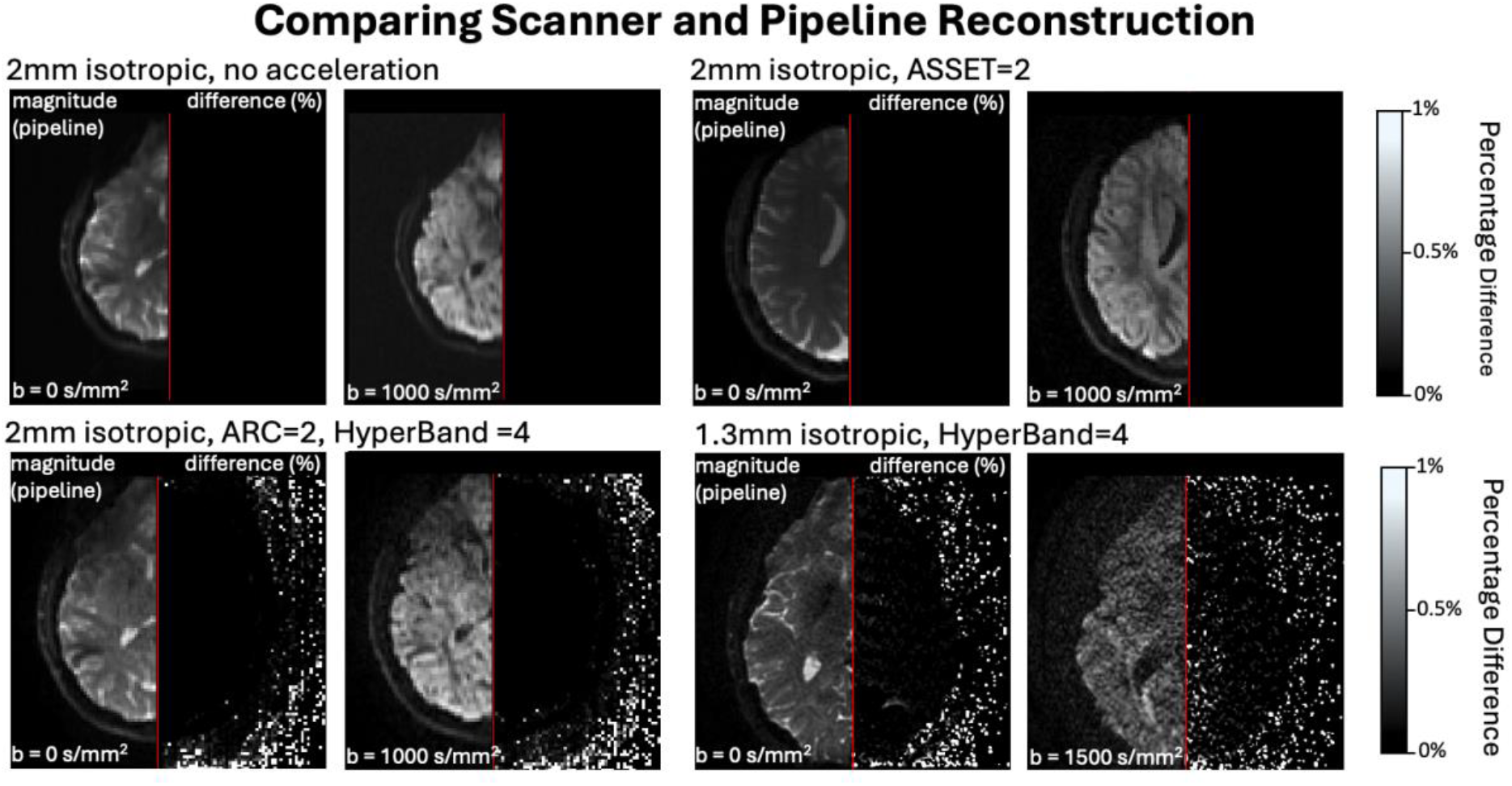
Comparison of offline magnitude and scanner reconstruction. Offline pipeline reconstruction and voxel-wise percentage difference in signal intensity between offline and scanner reconstruction for (top left to bottom right) 2mm isotropic no acceleration, ASSET = 2, ARC=2 + HyperBand=4 and 1.3mm isotropic HyperBand= 4 data. Minimal differences are observed when no acceleration or ASSET is used, and differences are below 1% in absolute value when HyperBand or ARC is used.

Figure 4a highlights some modifications made to the C++ Orchestra SDK to enable correct indexing of slices, correctly account for the reference scans in SMS acquisitions and correctly identify the phase encoding direction for blip-reversed acquisitions. We showcase examples of reconstructed complex images in Figure 4b, showing magnitude and phase reconstruction for a zero-filled acquisition and real-rotated homodyne acquisition (exemplified by negative values in the CSF signal). The offline pipeline also gives access to complex single-channel images and respective coil sensitivity profiles, as shown in Figure 4c, which can be retrospectively reconstructed using custom homodyne or POCS algorithms.

**Fig. 4.**
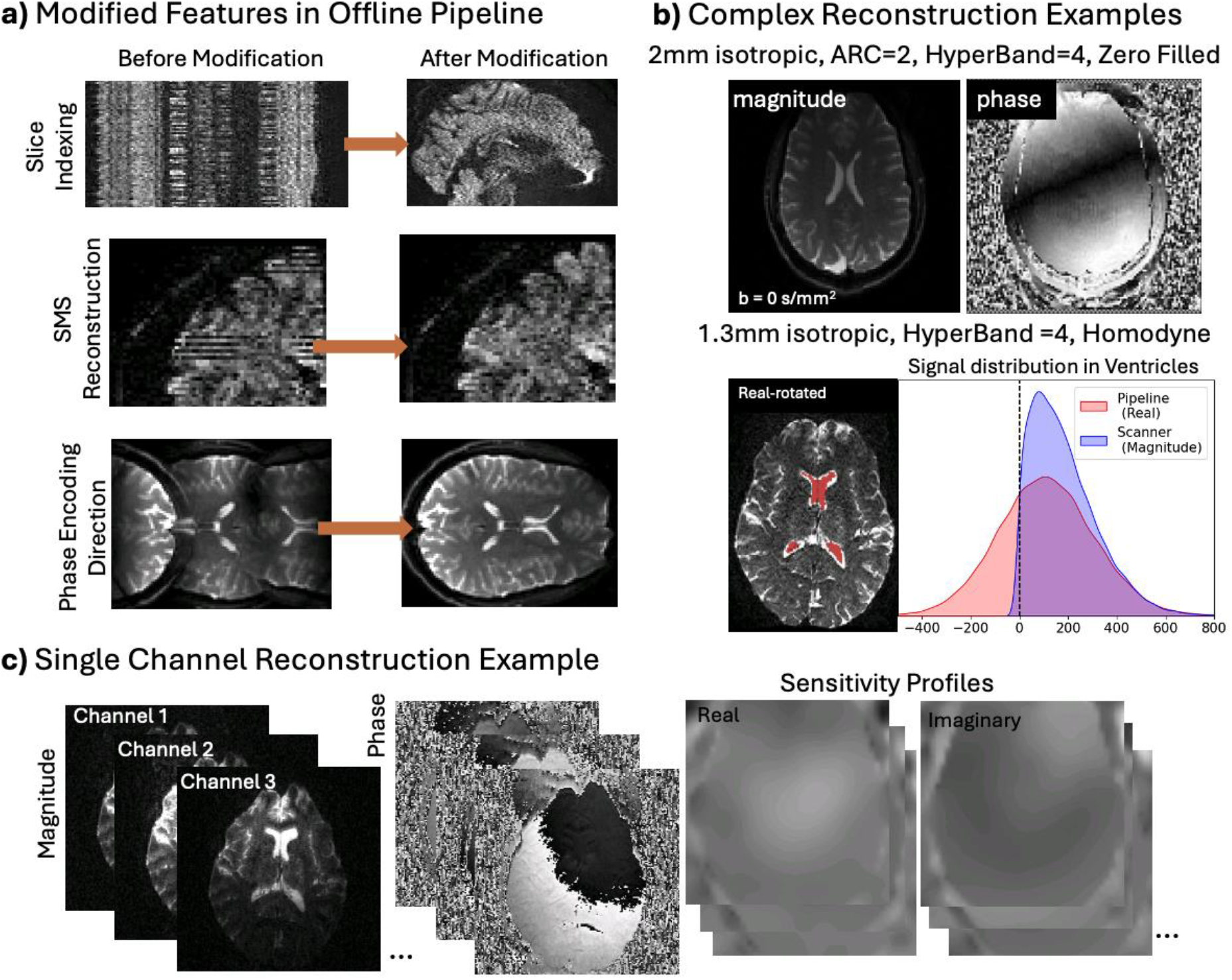
Offline reconstruction modifications and additional features. a) Images highlighting some modifications to the original Orchestra SDK reconstruction, from top to bottom: correcting slice indexing, SMS reconstruction and phase encoding direction reconstruction. b) Examples of pipeline complex zero-filled (magnitude and phase) or homodyne (real rotated) reconstruction, not available from scanner reconstruction. c) Example of single channel reconstruction showing single channel magnitude, phase and sensitivity profiles for channel combination.

### 3.2 Denoising Comparisons

The effect of denoising on exemplary slices for b=3000 s/mm^2^ (top) and b=1500 s/mm^2^ (bottom) shells is shown in Figure 5. To showcase the increased ability to discriminate anatomy post-denoising, sulci between the insula and adjacent lobes are highlighted. The location and size of the sulci is compared between high signal CSF in b=0 s/mm^2^ slices and low signal in diffusion weighted slices. All denoising approaches substantially improve the images over the raw data. MPPCA_SVS_ denoised diffusion slices show the highest anatomical correspondence with the b=0 s/mm^2^ slice. Noise floor-induced bias in magnitude denoised data (|MPPCA|_SVS_) is highly visible when compared to complex denoised data (MPPCA*_SVS_, NORDIC or ARDL), highlighting the advantages of denoising complex, pipeline-reconstructed data over magnitude-denoised data. Furthermore, as highlighted in the figure inlays, anatomical details are more visible when applying MPPCA*_SVS_ or NORDIC compared to ARDL, with MPPCA*_SVS_ showing qualitatively the best performance across all approaches.

**Fig. 5.**
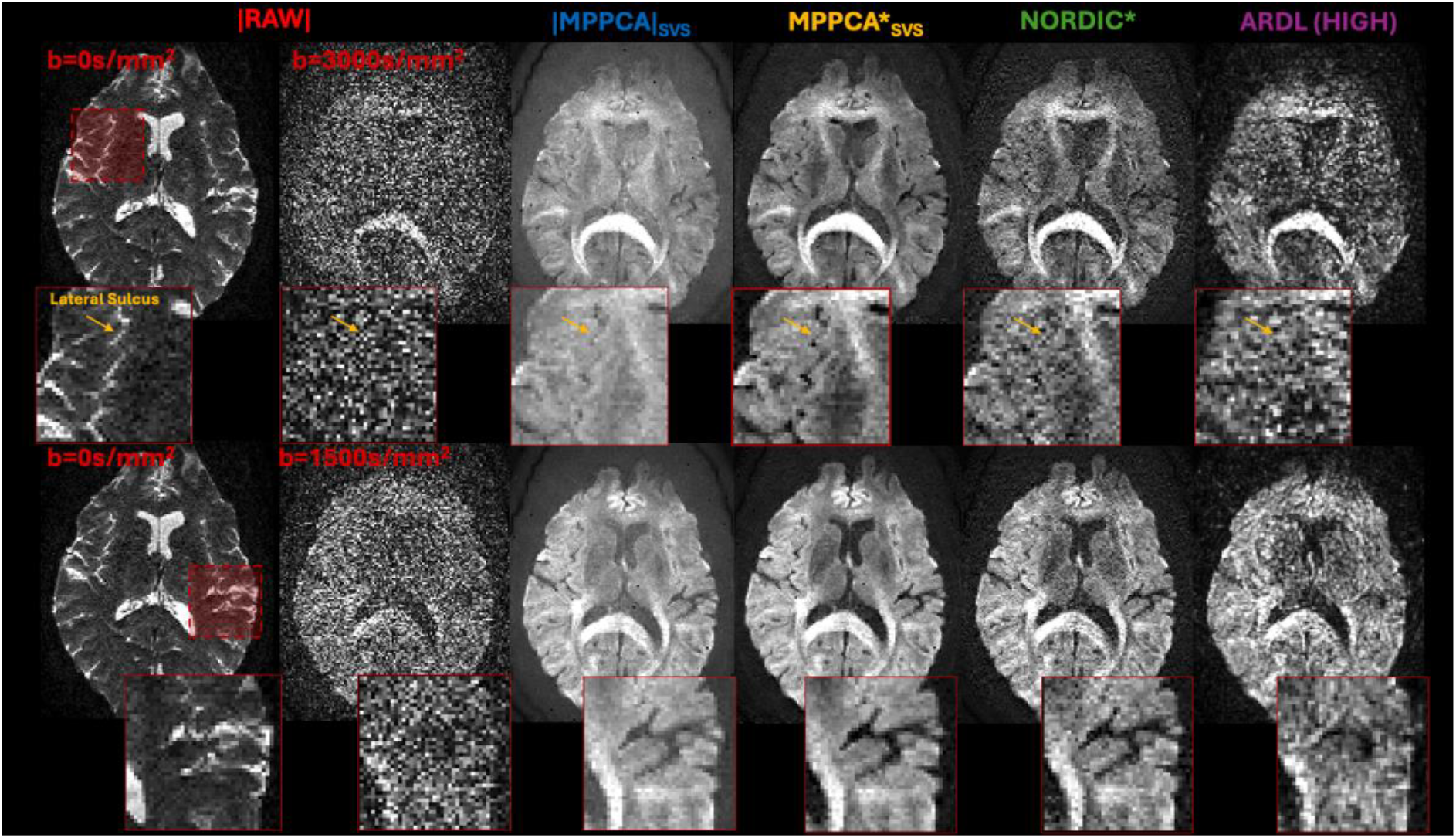
Comparison of raw and denoised data for b=3000 s/mm^2^ (top) and b=1500 s/mm^2^ (bottom) slices. A raw b=0 s/mm^2^ slice is shown on the left to provide anatomical benchmarks for diffusion weighted volumes. Different columns correspond to raw data, magnitude-denoised |MPPCA|_SVS_, complex-denoised MPPCA*_SVS_, complex-denoised NORDIC* and complex-denoised ARDL with high denoising weighting. Smaller panels highlight sulci (highly visible in b=0 s/mm^2^ raw data) to compare improvements in resolution and anatomical fidelity of denoised data.

We subsequently performed quantitative assessments. Figures 6–8 provide an overview of metrics related to changes in variance and bias of the signal due to denoising. Estimates of SNR across the b=0 *s*/mm^2^ volumes and angular CNR per shell are presented in Figure 6, showing better performance for 4D PCA-based denoising methods compared to 2D ARDL.

**Fig. 6.**
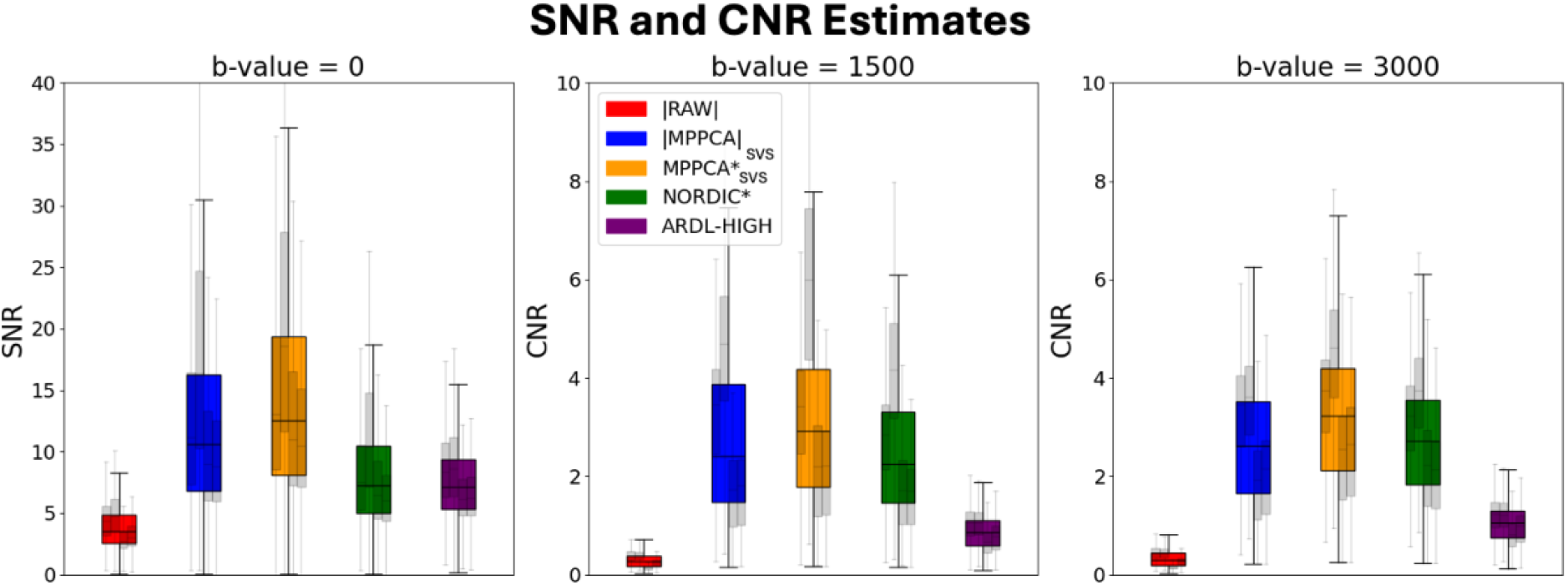
SNR (across b=0 s/mm^2^ volumes) and angular CNR estimates using FSL-EddyQC, thinner grey box plots represent single-subject data.

Figure 7a presents the dMRI signal attenuation at a given b value, having the signal reordered based on the dot product of the corresponding diffusion-sensitisation gradient direction and the principal DTI V_1_ in each voxel (i.e. parallel to perpendicular the main fiber orientation) and then averaged across a region of very anisotropic voxels (depicted by the mask in the body of the corpus callosum). We can observe how the signal is rectified in the magnitude domain and how all complex-domain denoising methods significantly increase the signal dynamic range and reduce the noise floor bias. The most significant decreases are associated with NORDIC and ARDL. In Figure 7b, the ADC peanuts present a similar picture with magnitude-domain denoising retaining a “squashed” peanut shape [7], which is recovered by all the complex denoising approaches.

**Fig. 7.**
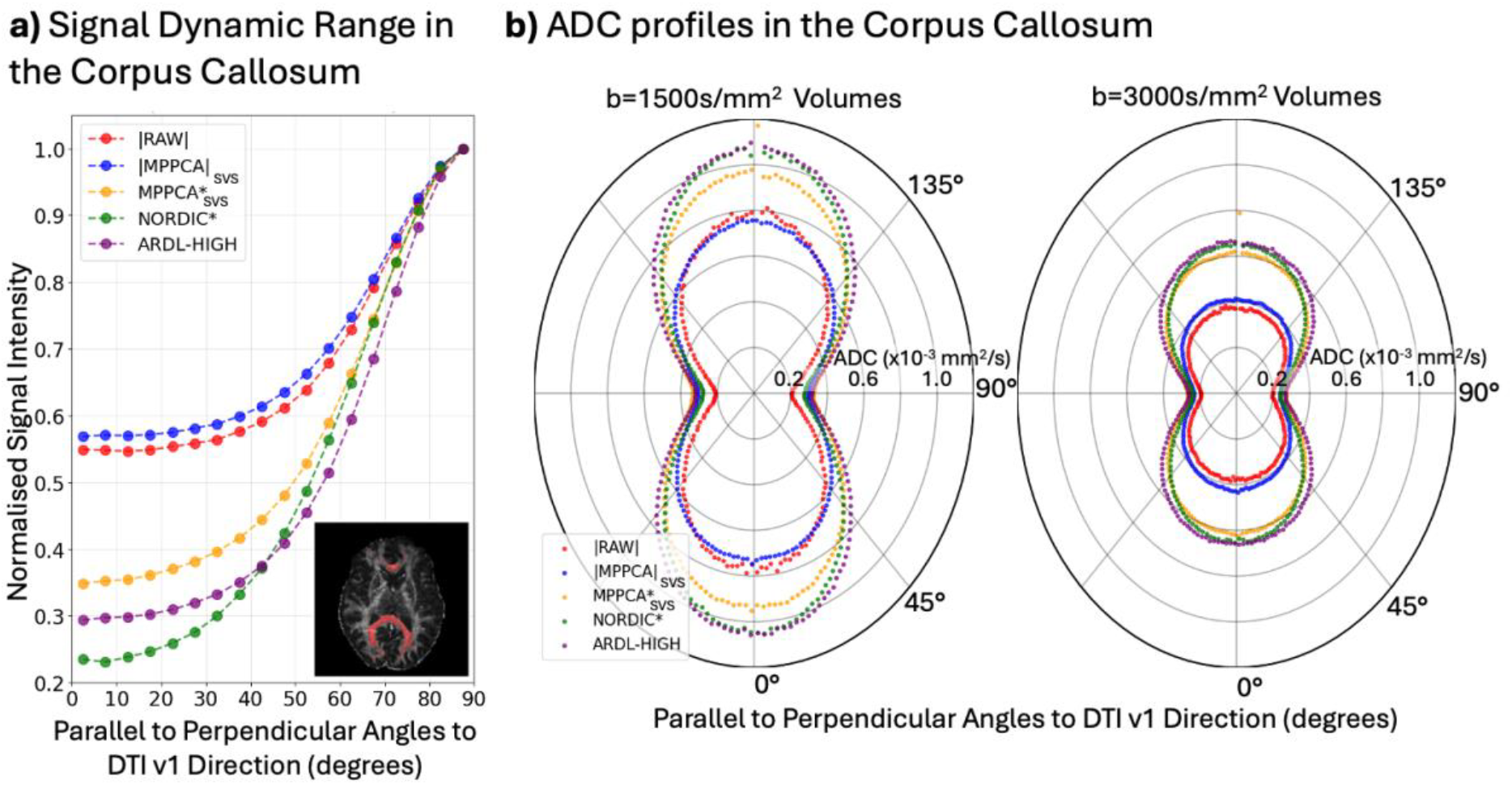
Signal dynamic range and ADC measurements in the corpus callosum obtained using a registered mask and an FA>0.6 mask from RAW data (panel highlights example high FA voxel mask in red). a) Signal attenuation is measured by binning b=3000 s/mm^2^ shell signals with respect to angle to DTI v1 direction and normalizing by the highest signal bin for each denoising mode. b) The ADC peanuts use the binned ADC measurement with respect to angle to DTI v1 direction for either the b=1500 s/mm^2^ or b=3000 s/mm^2^ shell.

Finally, Figure 8 presents the distribution of maximally attenuated CSF signal in the ventricles, a proxy for the noise distribution, again showing increased noise floor suppression when denoising in the complex domain, with ARDL having the best performance. Taken together, these results suggest that denoising in the complex domain reduces noise-induced bias and variance, with ARDL performing the best in the removal of bias in the ventricles, NORDIC performing best in the removal of bias in regions of high diffusion anisotropy, and MPPCA*_SVS_ performing the best in the reduction of variance.

**Fig. 8.**
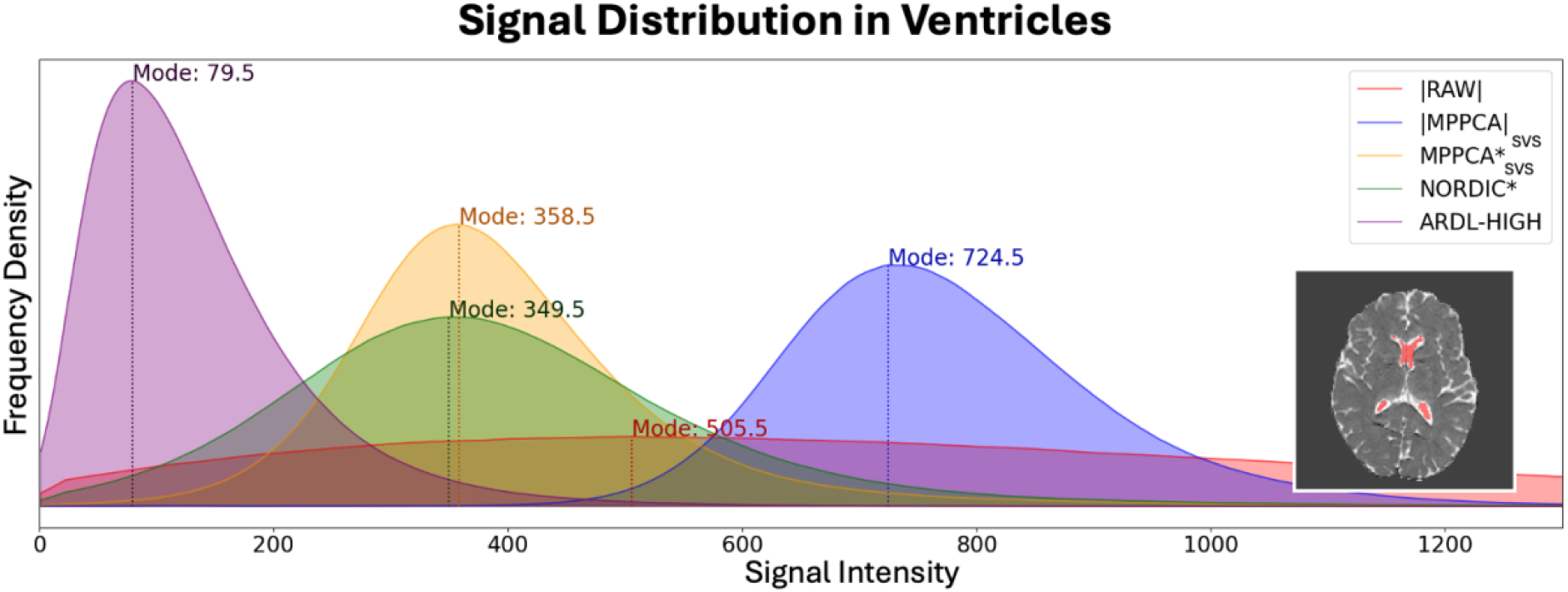
Distribution of signal in the ventricles as a proxy for the noise distribution. Inlay highlights example ventricle mask (red) for a single subject.

Downstream effects of denoising were assessed by exploring fiber orientation estimation and white matter bundle reconstructions, as seen in Figure 9. Fiber orientation estimations using the multi-shell ball & sticks model (up to 3 orientations per voxel) are shown in Figure 9a, highlighting estimates around the centrum semiovale, as a location of complex (up to three-way) fiber crossings. Both MPPCA*_SVS_ and ARDL denoised data show increased identification of two and three-way crossings, depicting orientations from the corpus callosum, the corticospinal tract and the longitudinal fasciculi running in and out of the coronal plane. These are mostly absent in the raw data. Also, MPPCA*_SVS_ estimates are more qualitatively spatially continuous than ARDL, implying better overall performance. It is important to note that FSL’s BedpostX uses an automatic relevance detection (ARD) algorithm [48] to select the number of crossing fibers in a voxel, using a data-driven approach to select between one and three fibers to model each voxel’s signal.

**Fig. 9.**
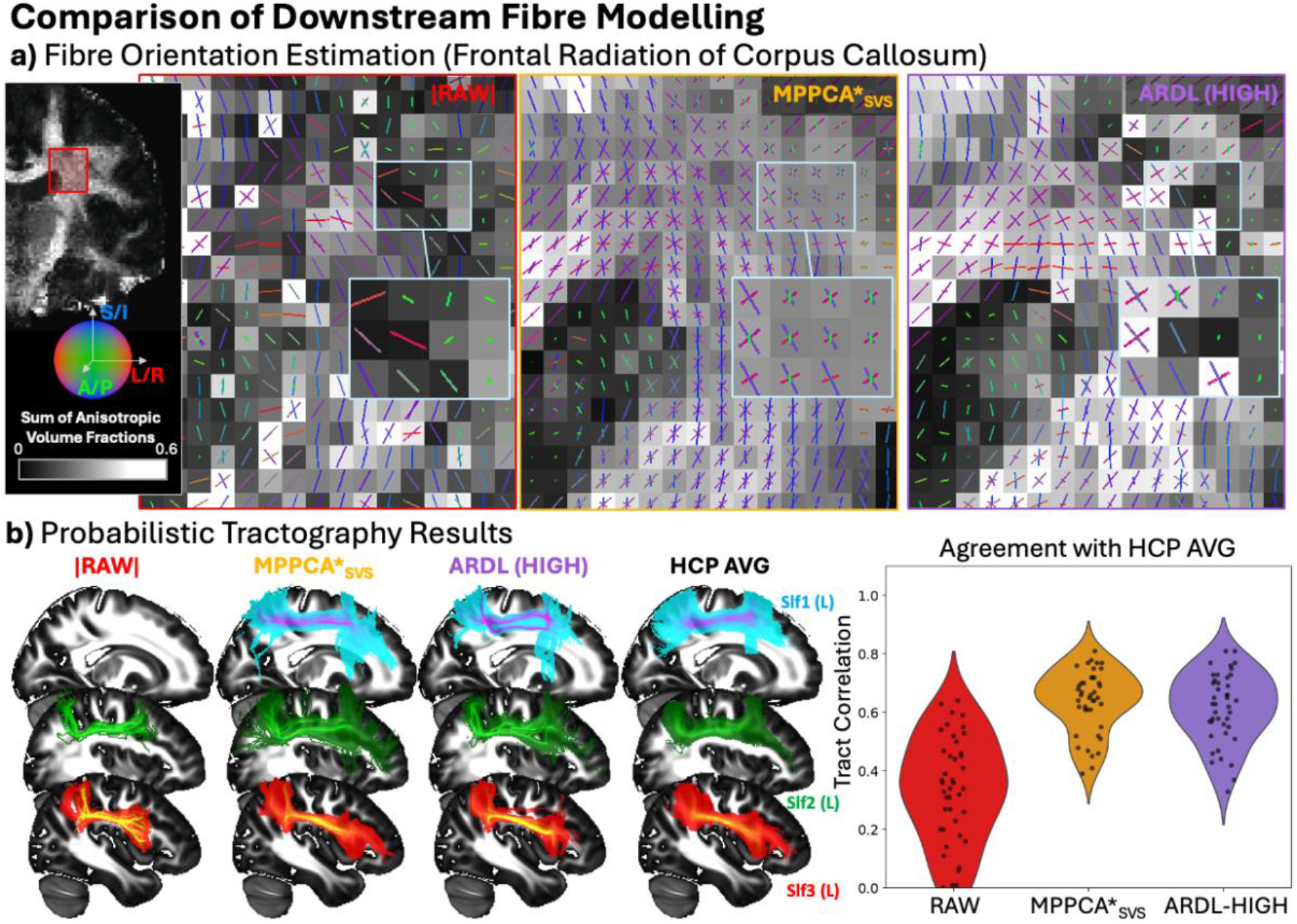
a) Comparison of fiber orientation estimation, using FSL’s BedpostX, highlighting regions of complex fiber architecture involving multiple crossing fibers. Sticks are length-scaled by their corresponding volume fraction. Voxel shade represents the sum of anisotropic volume fractions. b) Results from probabilistic tractography (average of 4 subjects) using FSL’s XTRACT showing maximum intensity projections of superior longitudinal fasciculi (left) and correspondence with a 50-subject HCP average for all 42 tracts (right). Path distributions were thresholded at 0.1% and correlated against the corresponding path distributions of the HCP average.

Fiber orientation estimates were subsequently used to perform probabilistic tractography and reconstruct 42 major white matter fiber bundles using FSL’s XTRACT. Figure 9b (left panel) examples participant-averaged tract density maps, highlighting results for the superior longitudinal fasciculi (SLF1, SLF2 and SLF3) and 50-subject HCP averages for comparison. We selected the SLFs as they pass through regions of complex fiber architecture. In the RAW data, tracking of the SLF1 was unsuccessful, whereas MPPCA*_SVS_ and ARDL successfully tracked the SLF1, though ARDL incorrectly fit streamlines which more likely belong to SLF2. Other smaller differences can be observed, for instance MPPCA*_SVS_ reconstructed SLF3 projects more anteriorly than the RAW or ARDL reconstruction. To quantify the differences further, violin plots on the right show the Pearson’s correlation between all 42 tract density maps of raw or denoised data and those of the HCP average, used as a reference. Denoising both with MPPCA*_SVS_ and ARDL significantly improved correspondence to the reference, with MPPCA*_SVS_ yielding better agreement overall.

## 4. Discussion

In this work, we presented a reconstruction pipeline for GE dMRI acquisition, which uses single-channel k-space data to give access to unfiltered, channel-combined complex data. The pipeline also gives the freedom to alter reconstruction steps like apodization filters, partial Fourier reconstruction method (POCS/Homodyne/Zero-filled), and gradient warp fields. In line with previous finding which used data from other manufacturers [1, 2, 5, 22], we demonstrate the benefits of denoising in the complex domain, using data obtained with the proposed pipeline, over magnitude-constrained denoising. This is particularly evident in reducing the effect of the noise floor bias, as observed in the signal dynamic range, ADC profiles in the corpus callosum, and noise distribution in the ventricles. Our offline reconstruction allowed us to obtain high-quality data at high spatial resolution and b value (1.3mm isotropic, b values up to 3000 s/mm^2^), similar to HCP dMRI [39], even if using a wide-bore scanner, such as the GE Premier, and a standard PGSE EPI sequence.

Previous work reports that denoising dMRI data reduces the variability of downstream metrics across scanners and protocols, with the greatest benefit when denoising in the complex domain [5]. So far, these comparisons have been limited to a single manufacturer across denoising domains [5], or using multiple manufacturers but constrained to magnitude domain denoising, with little control of the reconstruction algorithms used across vendors [49]. Our introduction of a flexible offline reconstruction pipeline for GE dMRI data allows to bridge the gap and allow cross-vendor and cross-domain comparisons in future work.

We showed that complex MPPCA_SVS_ performs better than NORDIC on these GE data in terms of contrast-to-noise gain and in the ability to resolve fine anatomical details post-denoising. This is different to previous results on comparing these two approaches with Siemens datasets [2]. These differences in trends are likely linked to the differences in MPPCA denoising methods used, with the MPPCA implementation from MRTrix (v3.0.7) [34] not including singular value shrinkage correction to compensate for noise-induced inflation of eigenvalues [21] or the option of patch averaging, as found in the DESIGNER implementation [35, 36] that we used in our study. Additionally, cross-vendor differences in partial Fourier reconstruction could also explain some of these differences in denoising outcomes [50]. Siemens reconstruction often uses zero-filled or POCS, whereas GE most commonly uses Homodyne reconstruction. The downstream denoising consequences of vendor-specific partial Fourier reconstruction methods are not well explored. However, even within a single vendor, these choices have been shown to significantly affect denoising outcomes [50].

We compared the performance of proprietary ARDL denoising and reconstruction, which benefits from within scanner integration and is modality agnostic, to open-source reconstruction and dMRI denoising using the offline reconstruction pipeline and commonly used PCA-based denoising methods [10, 11, 13]. These results suggest that PCA-based denoising methods outperform ARDL in terms of identifying fine image details, contrast to noise ratio gains, and fiber orientation estimation. This is not surprising, given that ARDL is 2D. Interestingly, ARDL significantly outperforms other methods when comparing the maximally suppressed CSF signal intensity in the ventricles, showing significantly lower noise-floor bias with a distribution closer to zero, but not a corresponding improvement in the signal dynamic range or ADC profile in regions of high diffusion anisotropy. This suggests that, post-denoising, the signal in ventricles, which is often used as a proxy for the noise distribution [2], does not capture the biophysical complexity to yield a generalizable metric of bias reduction across tissues. However, ARDL also operates in the complex domain and the overall benefits in reducing noise-induced biases agree with benefits offered by PCA-based denoising approaches when applied to complex data.

ARDL requires a 256×256 input matrix size which was achieved by embedding the acquisition matrix (160×160) into a zero-filled oversized matrix, followed by a post-ARDL spline interpolation to the acquisition matrix size for comparison against the open-source denoising methods. To ensure our trends were not driven by our custom post-ARDL interpolation, we carried out a comparison where all denoising methods were zero-filled to a 256×256 matrix size, using the same processing as the ARDL preprocessing, followed by the spline interpolation to the acquisition matrix size (160×160) prior to denoising. The trends in our results remained unchanged, as shown in Supplementary Figure S1. Additionally, the comparisons carried out in this paper only highlight the performance of ARDL on the ‘high’ setting, supplementary Figure S2 shows that this was the best performing mode available from scanner selection (low, medium, high). However, a limitation of this work is that we did not test ARDL capabilities beyond the ‘high’ setting, as ARDL uses a tuning hyperparameter ranging from 0 to 1, and the high setting corresponds to 0.75. Additionally, ARDL includes unringing features, which were not evaluated against open-source alternatives[51] in this study.

Previous work has suggested that denoising earlier in the reconstruction process is beneficial to denoising outcomes [1, 50], from the reduction of non-stationary g-factor contributions to noise [11, 52]. Although ARDL benefits from denoising single-channel complex data rather than channel-combined data, it is important to note that 2D ARDL does not utilize the full redundancy of 4D dMRI data, unlike the open-source PCA-based approaches, developed explicitly for use in dMRI data. The proposed pipeline provides access to single-channel complex data, enabling evaluations of denoising earlier in the reconstruction process and across partial Fourier methods.

## 5. Conclusion

Our offline reconstruction pipeline for GE dMRI acquisition was validated against scanner magnitude reconstruction and gives access to unfiltered complex domain data for acquisitions with in-plane and SMS acceleration. We found significant gains in dMRI data quality when using our offline reconstruction pipeline, allowing denoising in the complex domain, both for reducing noise-induced variance and bias. Open-source PCA-based (4D) methods outperformed proprietary 2D complex-domain denoising using deep learning (ARDL) in terms of qualitative resolution, reduction of noise-floor bias and variance.

## Acknowledgements

FD is supported by a studentship funded by GE HealthCare and the University of Nottingham. SW and SNS are supported by an ERC Consolidator Grant (101000969). We would like to also thank Tim Sprenger, Jaemin Shin, Gavin Houston and Suchandrima Banerjee from GE HealthCare for useful discussions. Additionally, we would like to thank Martin Graves and Ilse Patterson for giving us access to the GE Premier scanner at CUH, while the Nottingham one was out of action.

