## Supplementary Material for "Offline Reconstruction of Diffusion MRI Acquisitions for Comparison Between Complex PCA-based and AI-based Denoising"

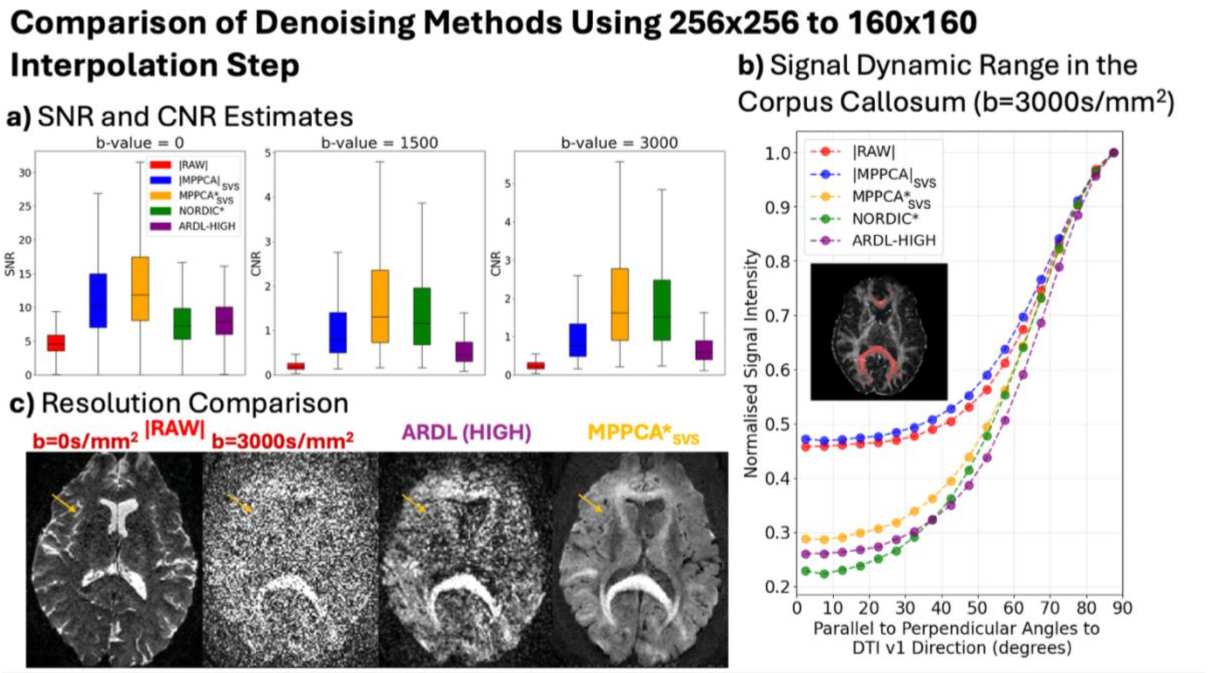


***Fig. S1.*** *Comparison of denoising outcomes with identical interpolation steps across methods using a single subject. a) SNR and CNR evaluation. b) Signal dynamic range in the corpus callosum. c) Comparison of resolution outcomes.*

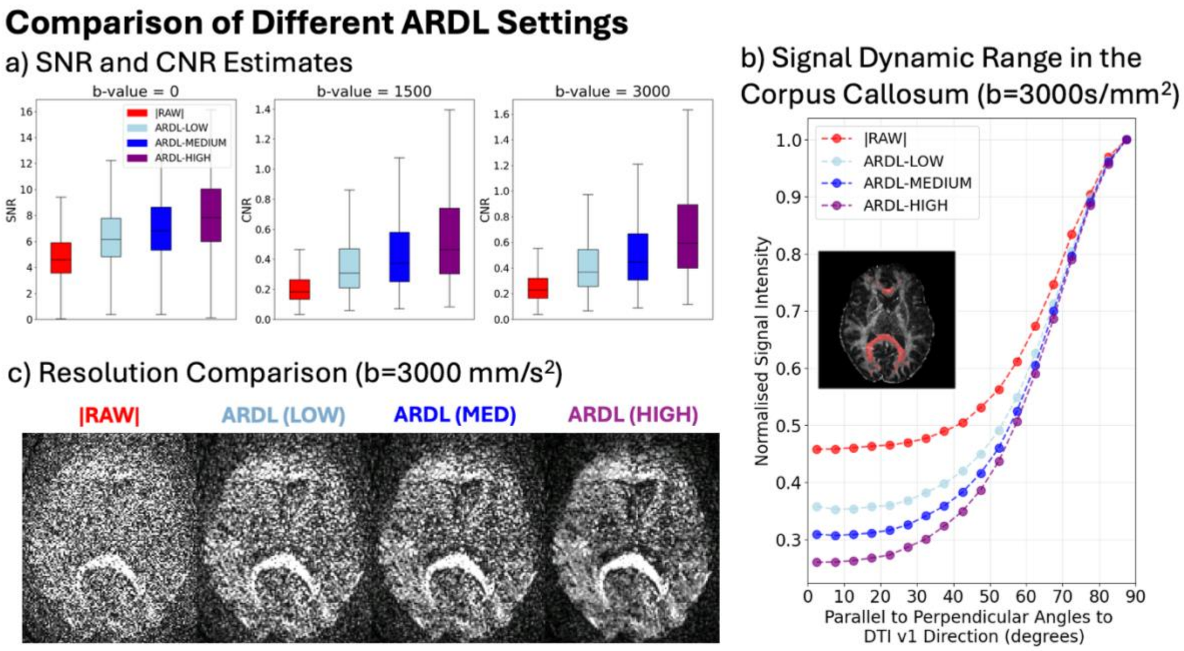


***Fig. S2.*** *Comparison of denoising outcomes across ARDL settings (low – 0.3, medium – 0.5, high – 0.75) using a single subject. a) SNR and CNR evaluation. b) Signal dynamic range in the corpus callosum. c) Comparison of resolution outcomes.*
